# Trans-kingdom delivery of aphid-derived small RNAs into *Arabidopsis thaliana* modulates plant immunity

**DOI:** 10.64898/2026.06.16.732591

**Authors:** Junlin Chen, Dimitrije Markovic, Claudia Cortes de Felipe, Elin Nygren, Maria Luz Annacondia, Velemir Ninkovic, German Martinez

## Abstract

Herbivore insects are on an evolutionary tug-of-war with plants. An important part of the plant-herbivore insect interaction is the exchange of molecules, in particular proteins (in the form of effectors) and RNA. Among RNAs, small RNAs have been identified as important molecules shaping the communication between different pests/parasites and their host, but the role of these molecules in the interaction between aphids and plants is not well understood. Here, we explored the role of aphid-derived sRNAs in the interaction between *Arabidopsis thaliana* and the green peach aphid, *Myzus persicae*. Using sRNA sequencing, we identified a significant amount of bona-fide aphid-derived sRNAs within Arabidopsis tissues. Using immunoprecipitation followed by sRNA sequencing and degradome sequencing we determined that these sRNAs are incorporated into endogenous AGO proteins, in particular AGO1, and induce the cleavage of transcripts involved in the modulation of the immune response against the aphid. Our results indicate that aphid sRNAs attenuate the immune response of *Arabidopsis thaliana* and can improve aphid performance. In addition, we identified that aphid-derived sRNAs are commonly injected into other aphid-host combinations. Accordingly, our work indicates that aphid-derived sRNAs are active players in aphid-host interactions.

## Introduction

Insects play vital roles in the stability of ecosystems, including pollination, decomposition of organic matter, regulation of other organisms via predation, and the dispersal of microorganisms between multiple hosts^1–4^. Approximately 50% of insect species are herbivores, and the majority of these show a pronounced host specialization^5^. Plant-insect interactions are mediated by multiple layers of recognition and response, including the reciprocal exchange of molecular cues at feeding interference^6^. Insect oral secretions contain herbivore-associated elicitors that are perceived by host plants and can trigger context-dependent responses in plants^6–9^. This view is further supported by the identification of both proteins and mRNAs that are directly delivered into host tissues through oral secretions^6–11^. Rather than being passive byproducts of feeding, insect-delivered molecules actively shape the host plant responses, facilitating herbivore feeding and establishment on the host^6–10^. Recent work has found that aphids deliver transcripts of unknown function into host plants^10^, suggesting a previously underestimated level of molecular interaction between aphids and their hosts.

RNA interference (RNAi) is an evolutionarily conserved sequence-specific gene regulatory mechanism mediated by small RNAs (sRNAs)^12,13^. Despite its conservation across eukaryotes, this mechanism has evolved in complexity experiencing specializations of its mode of action at the tissue and cellular levels. In plants, this mechanism, usually referred to as RNA silencing plays a central role in regulating gene expression and epigenetic modifications, which makes it crucial in the orchestration of development, maintenance of genome stability, and the response against stresses^12,14^. Two major branches of RNA silencing are generally recognized: transcriptional gene silencing (TGS), which mainly involves RNA-directed DNA methylation (RdDM) in plants, and post-transcriptional gene silencing (PTGS), which results in mRNA cleavage or translational repression^15,16^. Both pathways are initiated by double stranded RNAs (dsRNAs) of different origins, which are processed into sRNAs by DICER-LIKE (DCL) proteins. These sRNAs are subsequently loaded into ARGONAUTE (AGO) proteins and guided to complementary RNA or DNA sequences to exert their regulatory functions. The PTGS pathway is characterized by typical 21-/22-nt sRNAs that are usually loaded in AGO1/2/5/7/10, while the TGS pathway uses 24-nt sRNAs and AGO3/4/6/8/9^12,17^.

Unlike mammals, which have a highly evolved immune system, plants rely on RNA silencing as their primary defense against pathogen invasion, as the main antiviral mechanism^18^. Pathogen infection often alters hosts’ PTGS and TGS sRNA profiles, which in turn fine-tune immune responses^19^. A well-studied case is the regulation of Nucleotide-Binding Leucine-Rich Repeat (NB-LRR) resistance genes, whose rsiattack^20,21^. These examples illustrate that sRNAs function as crucial switches in plant immunity. Recent studies have revealed that sRNAs can also participate in trans-kingdom regulation between hosts and their interacting organisms (usually pathogens and parasites). Several examples of this exchange of sRNAs between different group species have been found, including the interactions between humans and bacteria or insects, and bacteria and *Caenorhabditis elegans*^22^. Interestingly, most of the examples of trans-kingdom RNA silencing have been reported in plants^23–31^. Importantly, several of these reports have identified that sRNA exchange between organisms is physiologically relevant. Plants can transfer sRNAs into their commensal organism to control its gene expression^26,27^ and, conversely, plant pathogens exploit sRNAs to silence host defense genes. Both bacteria and fungi deliver sRNAs into their hosts to suppress host immunity-related genes^27,28^. These discoveries established trans-kingdom RNA trafficking as an important mechanism of host–pathogen interactions.

Plants and insects have been reported to use this exchange of sRNAs to interact with each other. The monophagous planthopper *Nilaparvata lugens* secretes miRNAs into rice host plants to enhance its feeding^32^. In addition, the polyphagous aphid *Myzus persicae* is affected by *Arabidopsis thaliana*-secreted sRNAs^33^. However, whether sRNAs play a role in the host-pest interaction during aphid infestation remains largely unknown. Aphids are among the most destructive insect pests, causing significant crop yield losses worldwide, making the understanding of these mechanisms highly relevant for improving their control in the field. In this study, we investigated the role of aphid-derived sRNAs in the interaction between *Arabidopsis thaliana* and *M. persicae*. We identified abundant aphid sRNAs injected into *Arabidopsis* and loaded preferentially into AGO1, that have the ability to silence host mRNAs associated with plant immunity and enhance aphid feeding. We further confirmed this mechanism of sRNA exchange in another aphid, *Acyrthosiphon pisum,* suggesting a conservation of this mode of action across aphid-plant interactions. Our work indicates that aphid-derived sRNAs play an important role in modulating the interaction between aphids and their host plants.

## Methods and materials

### Plant material and aphid infestation

Col-0 or transgenic *A. thaliana* plants at the 4-rosette-leaf stage were infested with 40 wingless aphid nymphs per plant. To avoid cross-contamination, mock and infested pots were covered with air-permeable cages and kept under long-day conditions for 72 hours. The above-ground parts of the plants were collected, and residual aphids were carefully removed using a brush before the tissue was ground for RNA extraction.

### AGO Immunoprecipitation

Immunoprecipitation was performed using AGO1 and AGO2 antibodies (Agrisera), Dynabeads™ Protein A for Immunoprecipitation (Invitrogen), and IP buffer (20 mM Tris-HCl, pH 7.5; 5 mM MgCl□; 300 mM NaCl; 0.1% NP-40) supplemented with one tablet of Pierce Protease Inhibitor per 50 mL (Thermo), following the protocol of McCue et al. (2012). RNA was subsequently extracted using TRIzol reagent (Invitrogen) according to the manufacturer’s instructions.

### RNA extraction and libraries preparation

The tissue was ground in liquid nitrogen with morter and pestle. RNA extraction was performed using TRIzol reagent (Invitrogen) following the manufacture’s instruction. mRNAs were specifically isolated from the total RNAs using Poly(A) mRNA Magnetic Isolation Module (NEBNext), followed by the RNA-seq library construction using Ultra^™^ II Directional RNA Library Prep Kit for Illumina (NEBNext). sRNA-seq libraries were produced using Multiplex Small RNA Library Prep Set for Illumina (NEBNext) with gel-enriched sRNAs as described in Martinez et al. (2016). PARE-seq libraries were prepared as described by Zhai et al. (2014) using mRNA-enriched fractions obtained with the Poly(A) mRNA Magnetic Isolation Module (NEBNext). For all the libraries involved in this research, three bioreplicates were generated each contains 8-10 plants grown in two pots. Sequencing was conducted by Illumina^®^ using the NovaSeq platform for sRNA sequencing with single-end 50 bp reads, and the NovaSeq X Plus Series platform for mRNA sequencing with paired-end 150 bp reads.

### EGFP expression analysis (RT-qPCR)

Three bioreplicates of each transgenic line (Target#1, Target#2, Target#3) and treatment (mock and infested) were used for the expression analysis. RNA was extracted as previously described followed by the treatment of DNase I, RNase-free (ThermoFisher Scientific). cDNA was synthesized using the RevertAid First-Strand cDNA Synthesis Kit (ThermoFisher Scientific). 20 ul qPCR reactions were prepared using 5x HOT FIREPol EvaGreen qPCR Mix Plus (ROX) (Solis Biodyne) and conducted on a CFX Connect Real-Time PCR Detection System (BIO RAD). The Cq (cycle quantification) values got from the raw data were firstly normalized by the Cq value of ubiquitin coding gene, *ubq10*, then normalized by the Cq value of corresponding mock samples.

### Aphid settling analysis

Col0 and STTM transgenic plants (STTM#1, STTM#2, STTM#3) at the 4-rosette-leaf stage were used for the aphid settling analysis. One leaf of equivalent size and shape was selected from each plant and placed inside air-permeable polystyrene tube (diameter 1.5 cm, length 5 cm). The lower end of the tube was sealed with a plastic sponge containing a slit through which the leaf was inserted. Ten wingless aphids of *M. persicae* were placed inside the tube. The upper end of the tube was sealed with cotton. The number of aphids that settled on the leaf were recorded after 2 hours which is sufficient time for aphids to settle and reach the phloem^34^. The number of aphids settled per leave was counted and recorded as one bioreplicate. In total, 36-37 bioreplicates per group were used for the analysis.

### sRNA and RNA -seq data analysis

Total sRNA and AGO-IP sRNA libraries were first adapter-trimmed, and the resulting reads were aligned to the *A. thaliana* genome (TAIR10) using Bowtie v1.3.1^35^, allowing up to 2 mismatches, to obtain total sRNAs or aligned to other reference indexes for sRNA categorization. To accurately distinguish aphid-derived sRNAs, reads that did not map to TAIR10 were selected and subsequently aligned to the MP genome using Bowtie with 0 mismatches allowed. The sRNAs with lengths ranging from 18-28 were extracted for the size distribution analysis. RNA-seq libraries were adapter-trimmed then aligned to the TAIR10 genome using STAR v2.7.10b^36^, allowing no more than 10 multimapping positions. Read count matrices were then normalized using DESeq2^37^ to obtain differential expression data.

### Aphid-derived sRNA targets identification and degradome profiling

To accurately identify the target genes of MP-sRNA, another approach was attempted. Instead of mapping to TAIR10 first to filter the plant sRNAs, the sRNA-seq data were directly mapped to MP genome using Bowtie v1.3.1 with 0 mismatches allowed. The trimmed degradome data were first truncated keeping the first 20nt sequences only. The mapped sRNA reads and the truncated degradome reads were subjected into PARESnip2 for the prediction of the sRNA-mediated cleavage events. High-confidence target sites were determined based on stringent PARESnip criteria. To remove the possible false positive events, the target genes in the events that have significantly 2-fold higher degradome reads in Infested plants compared to the mock plants were selected as the candidates. To further identify the AGO1 mediated cleavage events, the sRNAs in the candidate events were mapped to the AGO1-IP sRNA-seq data using Bowtie v1.3.1 with 0 mismatch allowed.

For the PARE-seq coverage analysis, the trimmed and truncated degradome reads were aligned to TAIR10 reference transcript database using Bowtie2 v2.5.4 under the --very-sensitive end-to-end mode (seed length = 20, 0 mismatches allowed in the seed). To capture the exact cleavage sites, only the coordinate of the 5′ terminal nucleotide of each mapped read was extracted as the 5P signal and converted into a coverage format. For both mock and infested conditions, data from three replicates were pooled to calculate the mean and SD, which were subsequently visualized using line plots.

### Gene ontology analysis

Gene ontology (GO) analysis was performed using the GO Term Enrichment tool of The Arabidopsis Information Resource (TAIR). P-values were manually calculated based on the single-tailed hypergeometric test.

## Results

### sRNAs from the aphid *Myzus persicae* are transferred into Arabidopsis

We previously explored sRNA distribution in plants infested with *Myzus persicae* for 72-hrs^38^. During that analysis we identified that sRNAs from aphid-infested tissues did not follow the characteristic 80-90% mapping to the TAIR10 genome observed in mock samples (Sup. Table 1). Several examples of trans-kingdom transport of different molecules (both RNA and proteins) have been identified in the last years, including examples from insect-derived sRNAs in plants^32^. To understand the origin of this unmapped group of sRNAs we explored their potential origin from *M. persicae* by mapping these to its genome. Our analysis revealed that our libraries originating from infested samples contained *c.* 13% of aphid-derived sRNAs (Fig. 1a). By comparison, mock libraries only contained 0.3% of aphid-mapping sRNAs that we considered random mapping artifacts (Sup. Fig. 1a).

**Figure 1.**
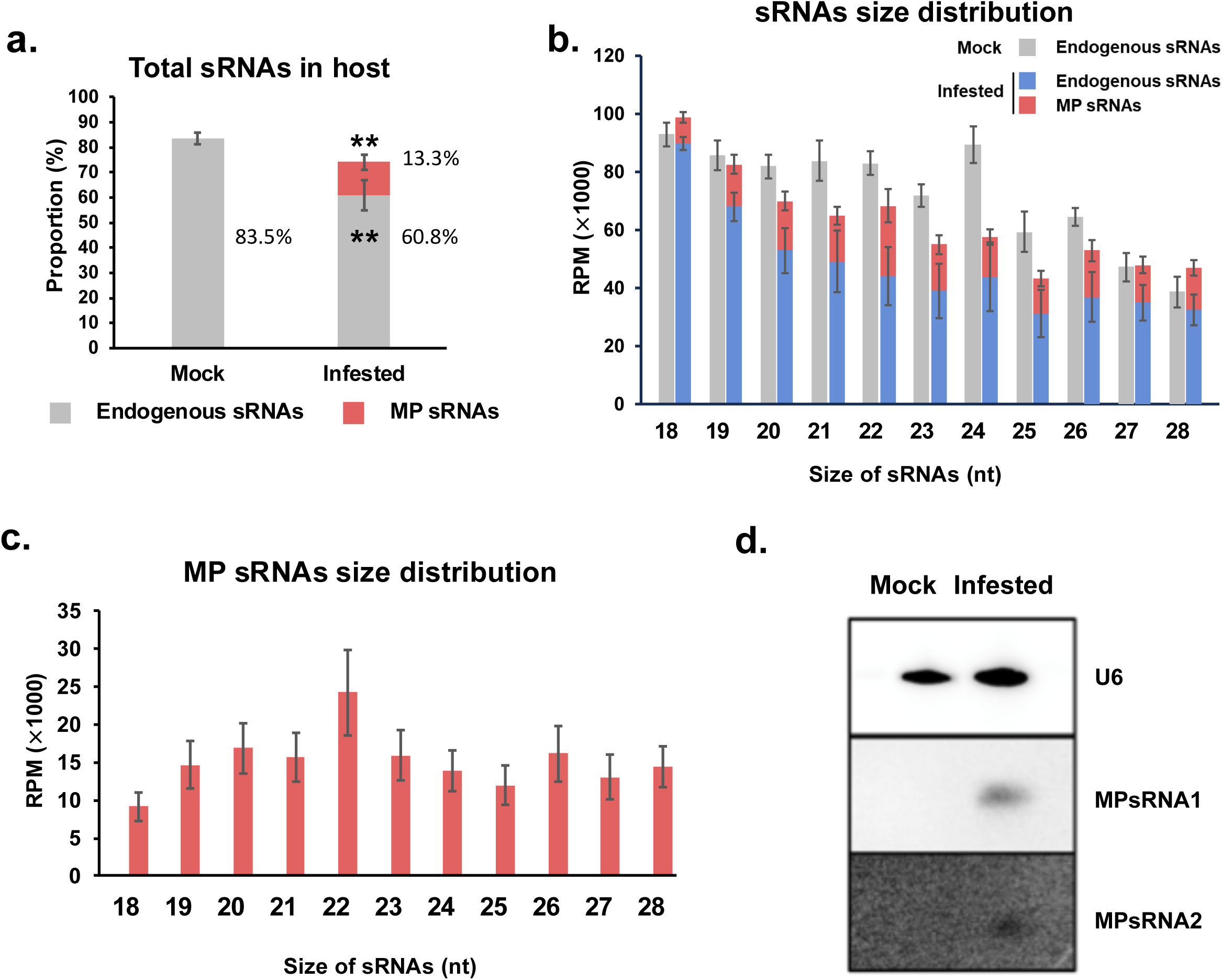
Bona-fide aphid-derived sRNAs are detected in host plants. **a.** Histogram showing the amount of sRNAs represented as percentage of the total sRNA libraries mapping to the *A. thaliana* TAIR10 genome (grey section of the bar) or to the *M. persicae* genome (red portion of the bar) in mock and infested samples. Error bars represent standard deviation of the mapping values obtained from 3 bioreplicates. ** = p-value<0.01. **b.** Histogram showing the size distribution of sRNAs in mock (grey color) and infested (blue+red color) libraries. For infested samples, plant (blue color) and insect (red color) sRNAs are represented. Error bars represent standard deviation of the mapping values obtained from 3 bioreplicates. **c.** Histogram showing the size distribution of *M. persicae*-mapping sRNAs (0 mismatches). Error bars represent standard deviation of the mapping values obtained from 3 bioreplicates. **d.** Northern blot of total RNA from mock and infested samples using probes against 2 highly accumulated aphid-derived sRNAs and U6 as a loading control.

Insect sRNAs differ significantly from plant sRNAs, with a predominant size of 22-nt, compared to the characteristic peaks of 21- and 24-nt of *Arabidopsis thaliana*^39^. To further analyze potential trans-kingdom aphid-derived sRNAs transferred into plants, we analyzed the size profile of these sRNAs. This analysis revealed that, as expected for bona-fide aphid-derived sRNAs, a majority of aphid-derived sRNAs were of 22-nt length, the characteristic length of insect sRNAs (Fig. 1b-c). We further confirmed the presence of aphid-derived sRNAs directly by Northern blot (Fig. 1d), which confirmed our sequencing results. Overall, our observation indicated that insect-derived sRNAs processed by the insect RNAi machinery were transferred into plants.

### Aphid-derived sRNAs are loaded into host AGO proteins

Next, we speculated whether aphid-derived sRNAs could integrate into the Arabidopsis endogenous RNA silencing pathways. We initially explored the alteration of endogenous sRNA categories by the presence of exogenous sRNAs (Fig. 2a). The presence of aphid sRNAs mainly affected miRNAs (1.7x decrease) and tasiRNAs (1.5x decrease) both reportedly loaded into AGO1^40,41^. Detailed analysis of miRNA profiles in mock and infested tissues supported this decrease observed in our categorization analysis, which showed that it significantly affected all miRNA (Fig. 2b) and tasiRNA (Sup. Fig. 1b) size categories. While other sRNA categories were also affected by the infection, these were reduced to a lesser extent (Sup. Fig. 1c).

**Figure 2.**
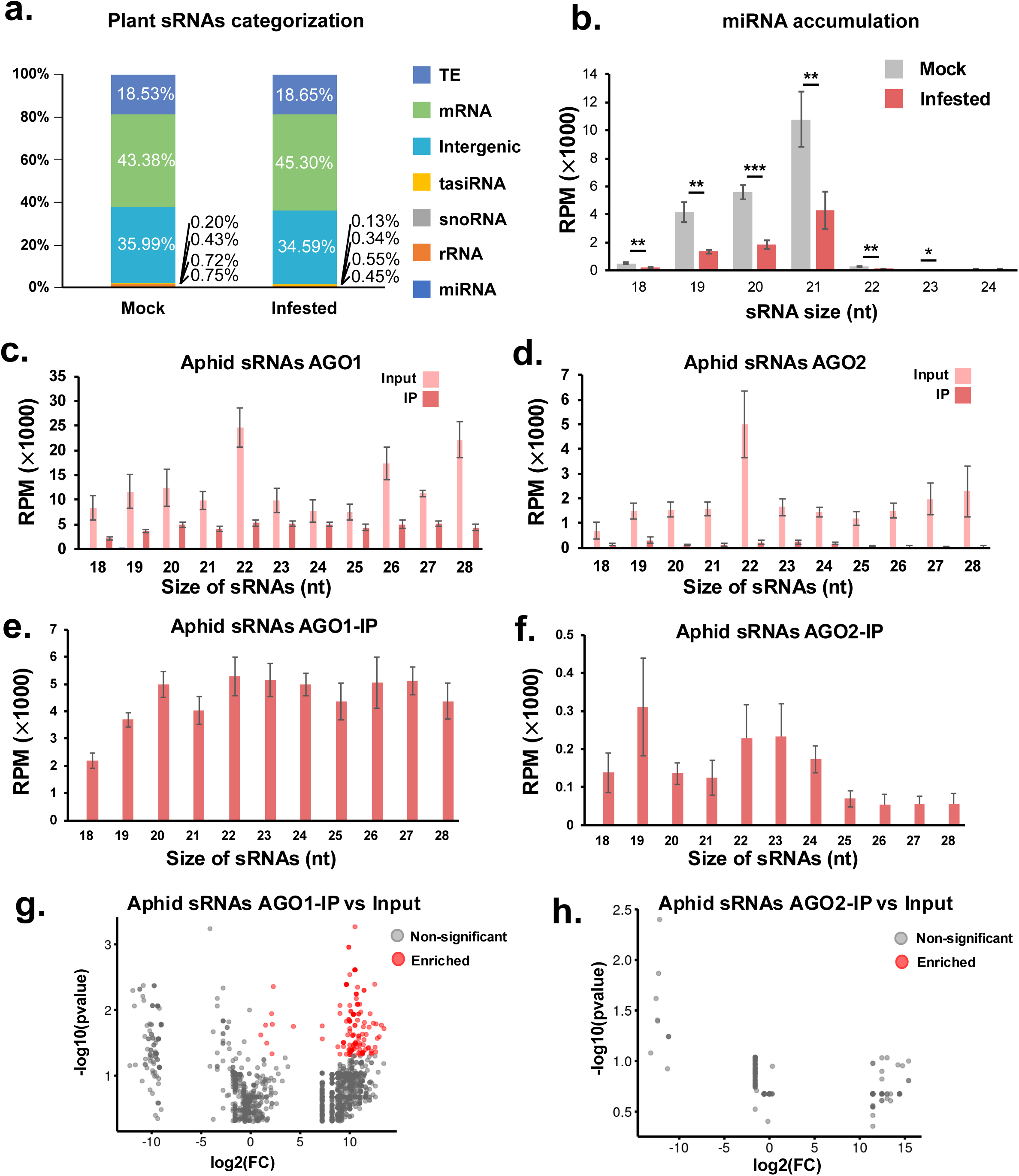
Aphid-derived sRNAs are loaded into AGO1. **a.** Categorization of sRNAs derived from mock and infested samples. Percentage of each category according to the total sRNAs mapped to the TAIR10 genome is indicated. **b.** Histogram showing the amount of miRNAs in mock and infested libraries. Error bars represent standard deviation of the mapping values obtained from 3 bioreplicates. **c-d.** Histograms showing the amount of sRNAs immunoprecipitated in AGO1 (c) and AGO2 (d) in input and immunoprecipitated fractions (IP) from infested libraries. Error bars represent standard deviation of the mapping values obtained from 3 bioreplicates. **e-f.** Histograms showing the amount of aphid-derived sRNAs immunoprecipitated in AGO1 (e) and AGO2 (f). Error bars represent standard deviation of the mapping values obtained from 3 bioreplicates. **g-h.** Volcano plots depicting the individual aphid-derived sRNAs significantly enriched (represented as red dots) in AGO1 (g) or AGO2 (h) IPs.

sRNA activity is mediated by their loading into AGO proteins, which determine their mode of action^42,43^. In somatic tissues, AGO1 and AGO2 monopolize the PTGS activity in plant-pathogen interactions^42^. Several pathogen/parasite-derived sRNAs have been reported to use the RNA silencing components of their host to exert different functions^44^. To directly test the incorporation of aphid-derived sRNAs into endogenous sRNAs, we analyzed their presence in AGO1 and AGO2 immunoprecipitated (IPed) fractions. Despite lacking evidence for AGO2 affection by aphid presence, we speculated that since this AGO protein is activated upon other biotic stresses such as viruses^45^, it might serve a similar role under insect infestations. As expected, AGO1- and AGO2-IPed sRNA categorization showed an enrichment in the miRNA category (Sup. Fig. 2a) and the expected enrichment in U and A 5’ nucleotides respectively (Sup. Fig. 2b), indicating the validity of the IPs. Next, we compared the amount of aphid sRNAs in both the input and IPs samples for both AGOs. Our initial analysis indicated that aphid-derived sRNAs in general were not particularly incorporated into any of the two AGO proteins (Fig. 2c-d). Despite this, aphid-derived sRNAs slightly maintained a preferential size of 22-nt in AGO1 and, partially, in AGO2 (were aphid-derived sRNAs of 19-nt were preferentially incorporated, Fig. 2e-f) although this enrichment was not significant. Detailed analysis of individual sRNA enrichment for both IPed fractions offered a different perspective, showing that 160 aphid-derived sRNAs were indeed enriched exclusively in AGO1 (Fig. 2g-h, Sup. Table 2). Interestingly, aphid-derived sRNAs accumulate to high levels in the input fractions of both AGO-IPs (Fig. 2c-d). This fact might reflect an overload of cell cytoplasm or apoplast with exogenous sRNAs with only a small, selected fraction being loaded into AGO proteins. Our analysis indicated that, despite being a relatively small part of the overall AGO-IPed fractions (Fig. 2a), aphid-derived sRNAs were loaded in AGO1.

### Aphid-injected sRNAs are active in Arabidopsis and modulate aphid feeding behavior

The transfer of aphid-derived sRNAs into host plants and their loading into host AGO proteins strongly suggested that these molecules could be active in the host tissues. We hypothesize that since these exogenous sRNAs were loaded into AGO1, they would exert the degradation of their targets. To address this question, we explored the function of the three most abundant aphid-derived sRNAs detected in host tissues. Using the sequences of those candidate sRNAs, we generated transgenic plants carrying fully complementary sequences fused to a GFP reporter to facilitate the tracking of the transgenic mRNA produced (Fig. 3a). Next, we simulated the conditions on which aphid-derived sRNAs were initially identified which involved the infestation of the transgenic lines with 40 adult aphids for 72 hrs before tissues sampling^38^. Analysis by RT-qPCR of GFP mRNA levels in the transgenic lines compared to their mock counterparts (same experimental conditions without the presence of aphids), showed that the expression levels of GFP were significantly reduced in all three transgenic lines (Fig. 3b). This result suggested that these artificial targets are recognized and silenced by aphid-derived sRNAs.

**Figure 3.**
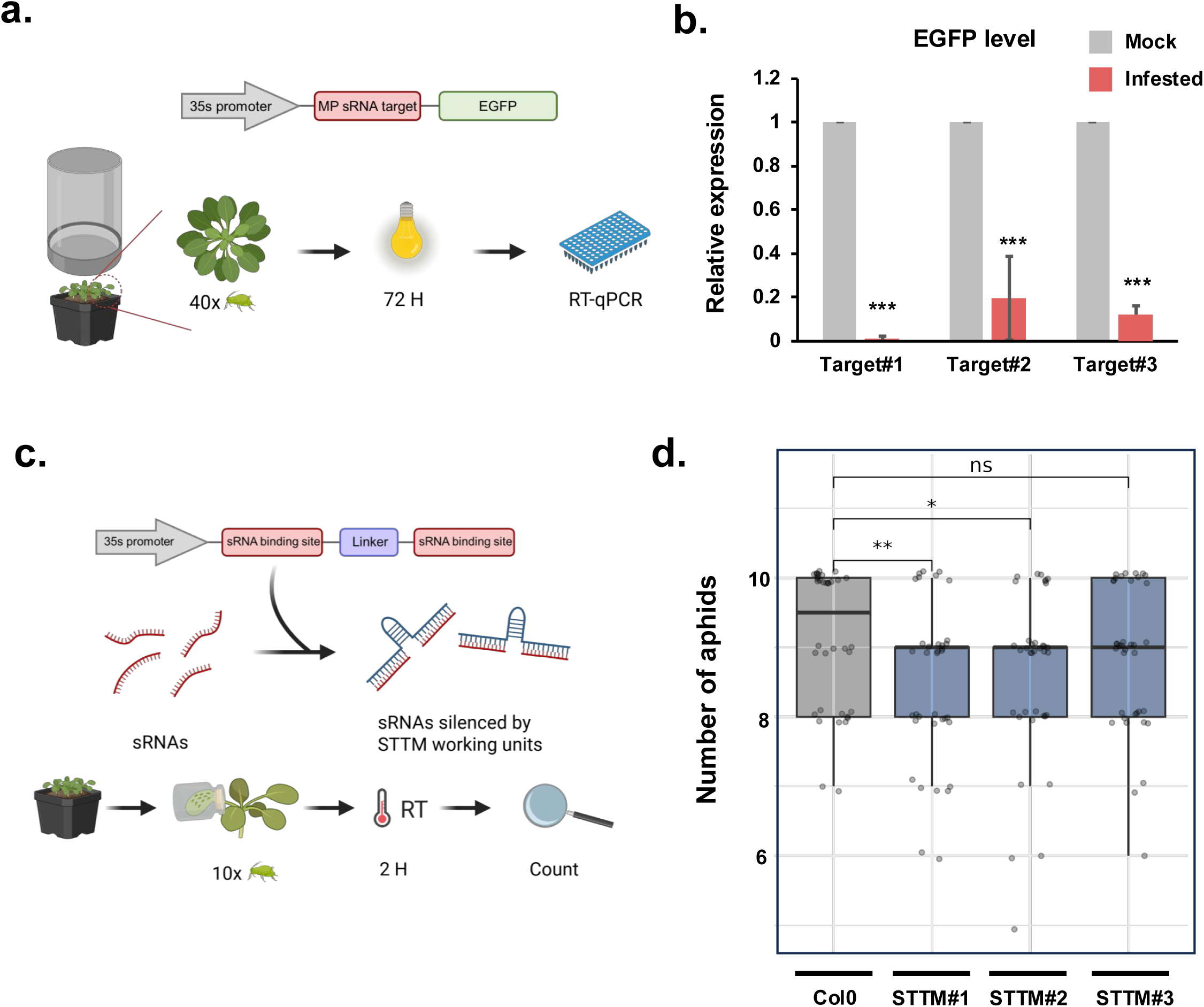
sRNAs delivered by aphids can mediate PTGS of endogenous transcripts and affect aphid behavior. **a.** Schematic representation of the transgenic lines generated to explore aphid-derived sRNAs PTGS activity, and the experimental set up to test their effect. In brief, 40 aphids were caged with each transgenic plant for 72 hrs. After this, aphids were manually removed and total RNA was extracted from the plants. **b.** qRT-PCR for eGFP expression in mock (grey bars) and infested (red bars) transgenic lines. Error bars represent standard deviation of the mapping values obtained from 3 bioreplicates. ***= p-value<0.001. **c.** Schematic representation of the transgenic lines generated to explore aphid-derived sRNAs physiological effect, and the experimental set up of the aphid settling test. In brief, 10 aphids were caged with a leave from each transgenic plant for 2 hrs. After this, aphids actively feeding from the leaves were inspected and recorded. **d.** Box-plot representing the amount of aphids actively feeding after 2 h in the indicated lines. *= p-value<0.05, **=p-value<0.01. n.s.=not significant.

Following the observation of their biological activity, we wondered if these exogenous sRNAs could contribute to aphid virulence. To further assess the functional relevance of these three aphid-derived sRNAs, we generated Short Tandem Target Mimic (STTM) transgenic lines with the aim to inhibit the activity of each candidate sRNA (Fig. 3c). We then subjected the generated STTM transgenic lines to an aphid settling test, where 10 aphids were introduced to a caged leaf for 2 hours. After this time, aphid plant acceptance was evaluated by visual observation (Fig. 3c). This experiment showed a significant decrease in the number of aphids settled on plants in two of the STTM transgenic lines compared to their wt control (Fig. 3d). These results suggest that the presence of aphid-derived sRNAs contributed to aphid virulence by potentially modulating the plant response to stress, as impairing their activity in the host compromises aphid performance.

### Genome-wide analysis of aphid-derived sRNA-targeted plant transcripts indicates a role in the modulation of the host response

Following the identification of a potential role in mediating insect virulence, we aimed to precisely identify the host mRNAs targeted by aphid-derived sRNAs. To this aim, we generated and analyzed parallel analysis of RNA ends sequencing (PARE-seq) in mock and aphid-infested plants using the same tissue previously used for the generation of sRNA libraries. To identify target mRNAs potentially cleaved by these exogenous sRNAs we integrated PARE-seq, aphid-derived sRNA, and the Arabidopsis transcriptome using PAREsnip^46^. In addition, we refined this strategy by selecting target mRNAs showing an increase of 2-fold in the number of reads present at the predicted cleavage site in infested samples compared to mock samples. Following this strategy, degradome profiling of the predicted aphid sRNA targets revealed a clear accumulation of cleavage signals within a −25 to +25 nt window surrounding the predicted cleavage sites in aphid-infested plants, whereas such enrichment was not observed in mock-treated controls (Fig. 4a). This pattern indicates a significant increase in site-specific cleavage events at aphid sRNA target loci during infestation. Furthermore, these targets showed a significant increase in the number of PARE-seq reads (Fig. 4b), pointing to a more intense degradation mediated by the presence of exogenous sRNAs. In total, we identified 207 aphid-derived sRNAs that targeted 131 host transcripts (Sup. Fig. 3a and Sup. Table 3). Gene Ontology (GO) analysis of targeted genes revealed a strong enrichment of stress response–related categories (Fig. 4c), indicating a targeting of aphid-derived sRNAs of stress signaling pathways and the regulation of the immune response.

**Figure 4.**
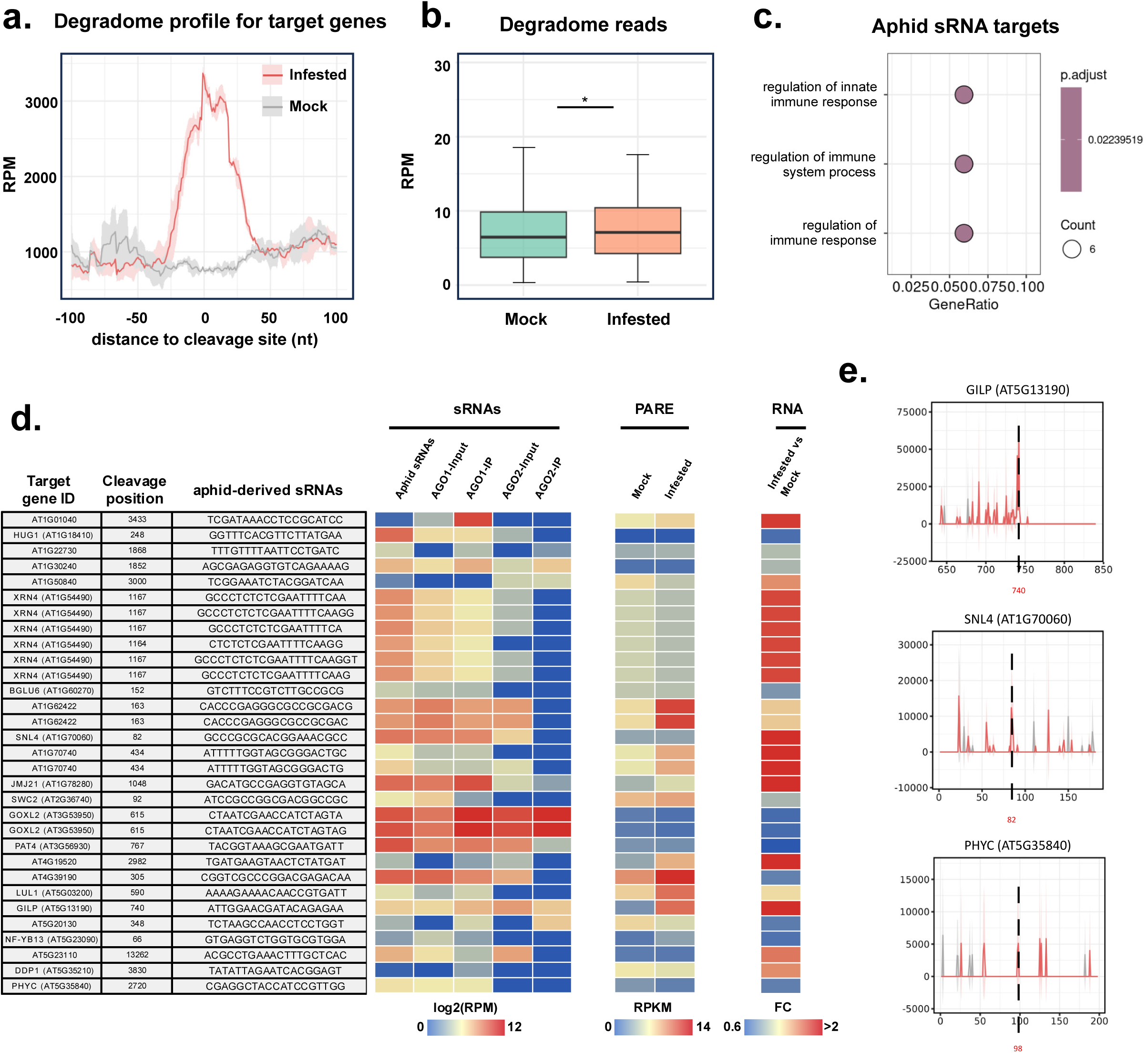
PARE-sequencing identifies endogenous transcripts targeted by aphid-derived sRNAs. **a.** Cumulative degradome profile coverage for mock (grey line) and infested (red line). 0 represents the predicted cleavage position identified by PAREsnip. **b.** Box-plot representing the amount of degradome reads in mock and infested samples. p-value is indicated. **c.** GO categories significantly enriched on identified targets of aphid-derived sRNAs. **d.** Table summarizing genes targeted by AGO1-enriched aphid-derived sRNAs. **e.** Cumulative degradome profile coverage in mock (grey line) and infested (red line) samples for 3 targets of AGO1-loaded aphid-derived sRNAs.

The identified target mRNAs included several cleaved by AGO1-enriched aphid sRNAs (Fig. 4d). These included important genes for plant immunity such as the negative regulator of cell death GILP^47^ and the reorganization of the transcriptional program such as SNL4^48^, both of which have been previously associated with the response to stress^49,50^ (Fig. 4d-e). Interestingly, despite being 22-nt and mediating the cleavage of their target mRNAs, aphid-derived sRNAs did not amplify the silencing cascade since we did not observe higher levels of 21- and 22-nt siRNAs derived from the targeted mRNAs (Sup. Fig. 3b). Nevertheless, mRNAs targeted by AGO1-loaded aphid-derived sRNAs produced a significant higher level of PARE reads, indicating that targeting by exogenous sRNAs increased their degradation (Sup. Fig. 3c). Collectively, our genome-wide analysis suggests that aphid-derived sRNAs mediate the cleavage of an important number of mRNAs that are important for the orchestration of the immune response elicited upon aphid infestation.

### Transmission of sRNAs is not an exclusive strategy of the *Myzus persicae-Arabidopsis thaliana* pathosystem

Next, we aimed to determine whether the aphid-delivered sRNAs from *M. persicae* represent a strategy unique from this polyphagous aphid or reflects a broader phenomenon among herbivorous insects. For this purpose, we reanalyzed publicly available sRNA sequencing data from different experiments. These included the interaction between the brown planthopper *Nilaparvata lugens* and rice (where trans-kingdom insect miRNAs were previously identified^32^), as well as datasets from the aphid species *Brachycaudus helichrysi* and *Aulacorthum solani* feeding on multiple host plant species including agriculturally important crop species (potato, alfalfa, and maize) and the wild species *Erigeon annua*, *Erigeon canadensis*, and *Solidago canadensis*^51^ (Sup. Table 4). In addition, we generated sRNA sequencing data for the interaction of the aphid *Acyrthosiphon pisum* in *Arabidopsis thaliana* at two different time points, 24- and 48-h post-infestation (Sup. table 1). Following the same sRNA mapping strategy as in our *M. persicae* analysis revealed that insect sRNAs were present in all these different combinations, ranging from 14% of the total sRNAs (interaction *B. helichrysi* and *A. solani-E. annua*) to 1% (interaction *N. lugens-*rice) (Fig. 5a). Importantly, insect-derived sRNAs were present in a 22-nt form, being this the major size identified in almost all samples (excluding the interaction between species mixture and *B. helichrysi* and *A. solani*) (Sup. Fig. 4a).

**Figure 5.**
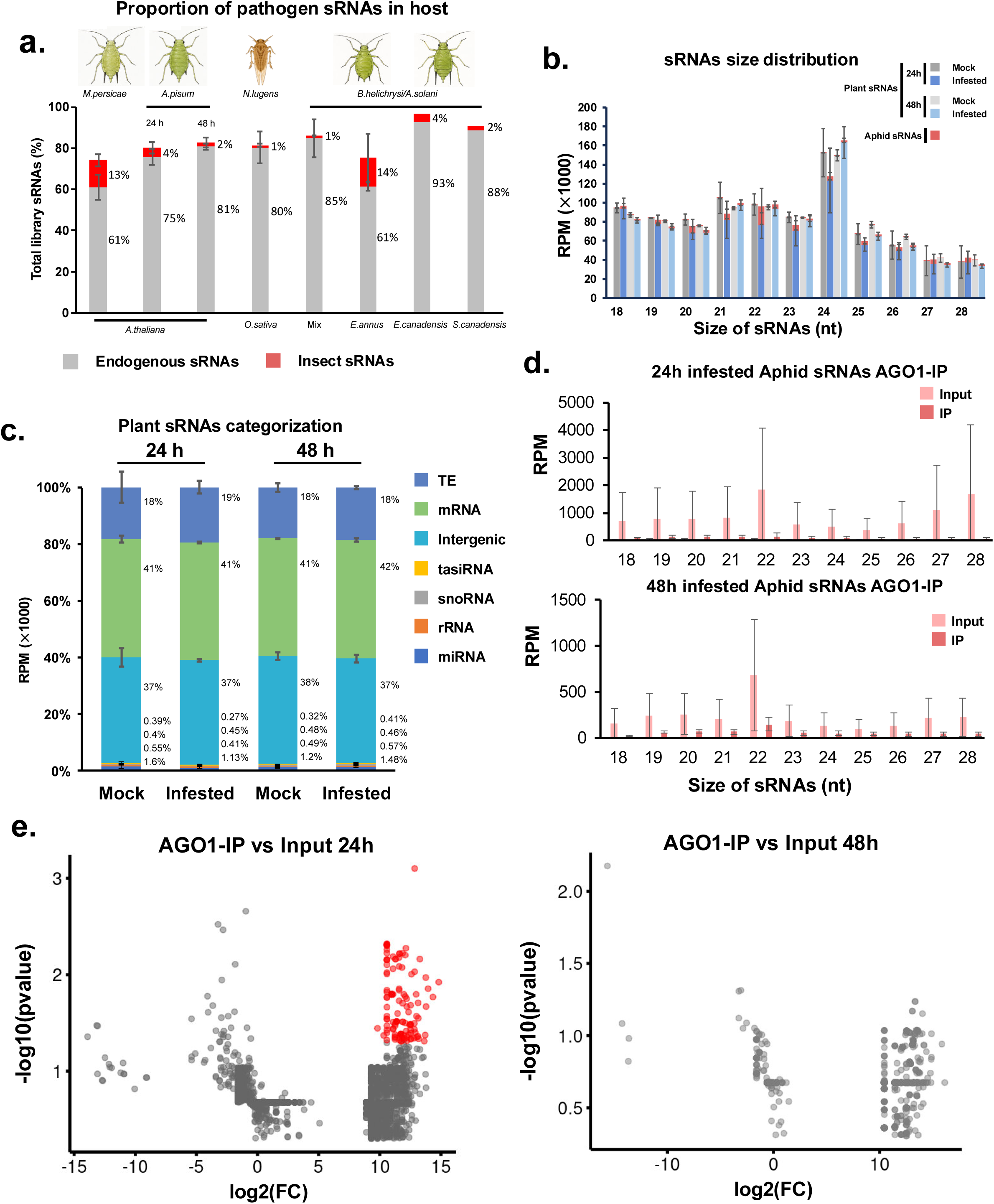
Injection of aphid-derived sRNAs is a common strategy among aphids in different hosts. **a.** Histogram showing the amount of sRNAs represented as percentage of the total sRNA libraries mapping to the indicated host plant (grey section of the bar, host name specie indicated in the bottom of the graph) or to the insect genome (red portion of the bar, name of the insect species indicated on the top of the graph). **b.** Histogram showing the size distribution of sRNAs in mock (grey color) and infested (blue+red color) libraries at the two time points analyzed, 24 and 48h post-infestation for *A. pisum*. For aphid infested samples, plant (blue color) and insect (red color) sRNAs are represented. Error bars represent standard deviation of the mapping values obtained from 3 bioreplicates. **c.** Categorization of sRNAs derived from mock and infested samples. Percentage of each category according to the total sRNAs mapped to the TAIR10 genome is indicated. **d.** Histograms showing the amount of sRNAs immunoprecipitated in AGO1 in input and immunoprecipitated fractions (IP) from infested libraries at 24 (top) and 48 h (bottom). Error bars represent standard deviation of the mapping values obtained from 3 bioreplicates. **e.** Volcano plots depicting the individual aphid-derived sRNAs significantly enriched (represented as red dots) in AGO1 IPs for 24 (left) and 48 h (right) samples.

To understand whether a similar interaction with the host RNA silencing pathways, in particular with AGO proteins, might take place in other insect-host interactions, we examined in detail the interaction between *A. pisum* and *A. thaliana*. Infestation with *A. pisum* resulted in a clear accumulation of 22-nt aphid-derived sRNAs in host tissues at the two time points under study (Fig. 5b and Sup. Fig. 4b). Categorization of sRNAs in *A. pisum*-infested plants revealed that, similar to our previous *M. persicae*-infested plant analysis, AGO1-loaded sRNA categories such as miRNAs and tasiRNAs were particularly affected by the insect feeding at 24h post-infestation (Fig. 5c, and Sup. Fig. 4 c-d). To determine if AGO1 was also loading these aphid-derived sRNAs we immunoprecipitated AGO1 fractions and generated sRNA libraries from these IPs at both time points. Similar to our previous AGO1 IP from *M. persicae*-infested samples, we also observed an enrichment of miRNAs, indicating the validity of the IP (Sup. Fig. 5a). Our AGO1-IP sRNA sequencing led to the identification of aphid-derived sRNAs at both time points, which conserved the characteristic 22-nt preferential size (Fig. 5d and Sup. Fig. 5b). Importantly, we identified 133 aphid-derived sRNAs significantly enriched into AGO1 only at 24h post-infestation (Fig. 5e and Sup. Table 5), pointing to a temporal dynamism in the accumulation of these sRNAs. In summary, our analysis revealed that injection of sRNAs into their host is a common strategy used by different insects colonizing diverse hosts, and that the interaction with host RNA silencing pathways (in particular AGO1) seems to be a common strategy for some of these insects.

## Discussion

The interplay between plants and insects represents a fascinating example of organism interactions with profound ecological relevance. Accumulating evidence suggest that plants and insects have reciprocally influenced each other, leading to events of co-evolution^52^. These events are the consequence of a multifaceted molecular arms race that involves the exchange of proteins and RNAs^6^. Recent works further indicate that multiple organisms (including insects) use sRNAs as signals mediating trans-kingdom communication^23–25^. Adding to that knowledge, our work shows that aphids (both *M. persicae* and *A. pisum*) use sRNAs as a trans-kingdom molecule that modulates the transcriptional response of their host plant, *Arabidopsis thaliana*. Interestingly, our work further indicates that sRNAs mediating host manipulation may extend beyond these model interactions, encompassing other aphid-host interactions such as the ones encompasing *B.helichrysi* and *A.solani* feeding on multiple host species including *E.annua, E.canadensis, S.canadensis, M.sativa, Z.mays,* and *S.tuberosum*. These results point to sRNA-mediated host manipulation as a conserved strategy among aphids to modulate their host plant response.

Our analysis of aphid-derived sRNAs indicate that these molecules are likely generated by aphids, since their prevalent size in most of the interactions analyzed here is 22-nts the characteristic, which is the main size of insect-produced sRNAs^39^. Aphids are well-known vectors of different RNA viruses^53,54^ and can also deliver endogenous mRNAs into their host plants^10^. Transport of viral particles into host plants is believed to take place via vesicles generated within the digestive system of the insect^55^. Recent works indicate that vesicles play an important role in the trans-kingdom exchange of sRNAs between hosts and pathogens/pests^26,56–58^. We hypothesize that a similar situation might be taking place here since we identified a remarkable amount of aphid-derived sRNAs into our AGO1/2 input samples, pointing to an overload with aphid-derived sRNAs of the apoplastic and/or cytoplasmic space of the host cells (Figs. 2c-d and 5d). Alternatively, it is also plausible that the high amount of aphid-derived sRNAs in our input sRNA libraries could be a consequence of a concert activity of all AGOs, similar to the situation in viral-infected cells^44^. Indeed, we also observed some amount of aphid-derived sRNAs into AGO2, although none were enriched in this protein. Further analysis of the role of all AGOs during insect-host interaction will shed light into this aspect of insect-derived sRNA biology.

Importantly, we report that aphid-derived sRNAs are active within their host cells and that they influence insect performance. First, we observed that transgenic plants containing reporter proteins fused to a perfect complementary sequence to highly accumulating aphid-derived sRNAs were silenced upon infestation (Fig. 3a). Furthermore, removing those same aphid-derived sRNAs from the cytoplasmic pool via STTM constructs caused a reduction in aphid performance in 2 out of 3 of our cases of study (Fig. 3b). Since we observed that several aphid-derived sRNAs were enriched in AGO1 (Fig. 2g) and that regular amounts of AGO1-loaded sRNAs were affected on infested plants (Fig. 2a), we hypothesized that the interference of these exogenous sRNAs with the host took place via their incorporation in the RNA silencing pathways of the host plants. This assumption was further supported by previous works were different host-pathogen/parasite interactions have been explored demonstrating examples of exogenous sRNAs being loaded into endogenous RNA silencing pathways and targeting endogenous mRNAs^23–31^. To further explore this potential interaction at the genome-wide level, we used GMUCT sequencing, which has been extensively used in plant biology to identify miRNAs/siRNA-targeted mRNAs^59^. Our analysis identified a relatively high number of genes targeted by aphid-derived sRNAs that furthermore have a role in the regulation of the immune response. This also included genes targeted by AGO1-enriched aphid sRNAs such as GILP and SNL4, which might be involved in regulating different aspects associated with aphid infestation such as hypersensitive cell death or the reprogramming of transcription^47–50^. Although counterintuitive, it is plausible to speculate that the targeting of genes orchestrating the immune response is a logical result considering aphid biology. Indeed, proteins secreted from aphid salivary glands prevent cell death and promote cellular recovery^60,61^. In view of this, it is plausible that, like aphid effector proteins, aphid-derived sRNAs modulate the immune response of plants to facilitate aphid performance. This idea is further supported by our analysis of aphid performance on STTM lines (which removed aphid-derived sRNAs from host plants), which showed a significant decrease in actively feeding insects, and was suggested in previous works on the role of trans-kingdom insect-delivered sRNAs^32^.

In summary, our study advances the understanding of the diverse array of molecules involved in the arms race between insects and plants, pointing to aphid-derived sRNAs as important molecules in the modulation of the plant immune response to aphid attack. In addition, our work exemplifies the importance of trans-kingdom sRNAs on the interactions between insects and their host plants.

## Acknowledgements

This research was supported by grants from the Swedish Research Council (VR 2021-05023 and VR 2025-05873), and the Novo Nordisk Foundation (NNF25OC0106663) to GM. The data handling was enabled by resources provided by the Swedish National Infrastructure for Computing (SNIC) at UPPMAX partially funded by the Swedish Research Council through grant agreement no. 2018-05973.

## References

1 Schoonhoven, L. M. Insect-plant relationships: the whole is more than the sum of its parts. Entomologia Experimentalis Et Applicata 115, 5–6 (2005). 10.1111/j.1570-7458.2005.00302.x

2 Chapman, R. F. The Insects: Structure and Function. 5 edn, (Cambridge University Press, 2012).

3 Holt, J. R., de Oliveira, N. C., Medina, R. F., Malacrinò, A. & Lindsey, A. R. I. Insect-microbe interactions and their influence on organisms and ecosystems. Ecology and Evolution 14 (2024). ARTN e11699 10.1002/ece3.11699

4 Pirttilä, A. M. et al. Exchange of Microbiomes in Plant-Insect Herbivore Interactions. Mbio 14 (2023). 10.1128/mbio.03210-22

5 Speight, M. R., Hunter, M. D. & Watt, A. D. Ecology of insects : concepts and applications. 2nd edn, (Wiley-Blackwell, 2008).

6 Howe, G. A. & Jander, G. Plant immunity to insect herbivores. Annual Review of Plant Biology 59, 41–66 (2008). 10.1146/annurev.arplant.59.032607.092825

7 Bleau, J. R., Gaur, N., Fu, Y. & Bos, J. I. B. Unveiling the Slippery Secrets of Saliva: Effector Proteins of Phloem-Feeding Insects. Molecular Plant-Microbe Interactions 37, 211–219 (2024). 10.1094/Mpmi-10-23-0167-Fi

8 Guo, H. J. et al. An Aphid-Secreted Salivary Protease Activates Plant Defense in Phloem. Current Biology 30, 4826-+ (2020). 10.1016/j.cub.2020.09.020

9 Jaouannet, M., Morris, J. A., Hedley, P. E. & Bos, J. I. B. Characterization of Arabidopsis Transcriptional Responses to Different Aphid Species Reveals Genes that Contribute to Host Susceptibility and Non-host Resistance. Plos Pathogens 11 (2015). ARTN e1004918 10.1371/journal.ppat.1004918

10 Chen, Y. et al. An aphid RNA transcript migrates systemically within plants and is a virulence factor. Proc Natl Acad Sci U S A 117, 12763–12771 (2020). 10.1073/pnas.1918410117

11 Song, J., Bian, J. G., Xue, N., Xu, Y. X. & Wu, J. Q. Inter-species mRNA transfer among green peach aphids, dodder parasites, and cucumber host plants. Plant Diversity 44, 1–10 (2022). 10.1016/j.pld.2021.03.004

12 Borges, F. & Martienssen, R. A. The expanding world of small RNAs in plants. Nat Rev Mol Cell Biol 16, 727–741 (2015). 10.1038/nrm4085

13 Svoboda, P. Key Mechanistic Principles and Considerations Concerning RNA Interference. Frontiers in Plant Science 11 (2020). ARTN 1237 10.3389/fpls.2020.01237

14 Kryovrysanaki, N., James, A., Tselika, M., Bardani, E. & Kalantidis, K. RNA silencing pathways in plant development and defense. International Journal of Developmental Biology 66, 163–175 (2022). 10.1387/ijdb.210189kk

15 Pickford, A. S. & Cogoni, C. RNA-mediated gene silencing. Cellular and Molecular Life Sciences CMLS 60, 871–882 (2003). 10.1007/s00018-003-2245-2

16 Erdmann, R. M. & Picard, C. L. RNA-directed DNA Methylation. PLOS Genetics 16, e1009034 (2020). 10.1371/journal.pgen.1009034

17 Zhan, J. P. & Meyers, B. C. Plant Small RNAs: Their Biogenesis, Regulatory Roles, and Functions. Annual Review of Plant Biology 74, 21–51 (2023). 10.1146/annurev-arplant-070122-035226

18 Lopez-Gomollon, S. & Baulcombe, D. C. Roles of RNA silencing in viral and non-viral plant immunity and in the crosstalk between disease resistance systems. Nat Rev Mol Cell Bio 23, 645–662 (2022). 10.1038/s41580-022-00496-5

19 Ruiz-Ferrer, V. & Voinnet, O. Roles of Plant Small RNAs in Biotic Stress Responses. Annual Review of Plant Biology 60, 485–510 (2009). 10.1146/annurev.arplant.043008.092111

20 Liu, Y. L., Teng, C., Xia, R. & Meyers, B. C. PhasiRNAs in Plants: Their Biogenesis, Genic Sources, and Roles in Stress Responses, Development, and Reproduction. Plant Cell 32, 3059–3080 (2020). 10.1105/tpc.20.00335

21 Fei, Q. L., Xia, R. & Meyers, B. C. Phased, Secondary, Small Interfering RNAs in Posttranscriptional Regulatory Networks. Plant Cell 25, 2400–2415 (2013). 10.1105/tpc.113.114652

22 Knip, M., Constantin, M. E. & Thordal-Christensen, H. Trans-kingdom Cross-Talk: Small RNAs on the Move. Plos Genetics 10 (2014). ARTN e1004602 10.1371/journal.pgen.1004602

23 Hudzik, C., Hou, Y. N., Ma, W. B. & Axtell, M. J. Exchange of Small Regulatory RNAs between Plants and Their Pests. Plant Physiology 182, 51–62 (2020). 10.1104/pp.19.00931

24 Katiyar-Agarwal, S. & Jin, H. L. Role of Small RNAs in Host-Microbe Interactions. Annual Review of Phytopathology, Vol 48 48, 225–246 (2010). 10.1146/annurev-phyto-073009-114457

25 Huang, C. Y., Wang, H., Hu, P., Hamby, R. & Jin, H. L. Small RNAs - Big Players in Plant-Microbe Interactions. Cell Host & Microbe 26, 173–182 (2019). 10.1016/j.chom.2019.07.021

26 Cai, Q. et al. Plants send small RNAs in extracellular vesicles to fungal pathogen to silence virulence genes. Science 360, 1126–1129 (2018). 10.1126/science.aar4142

27 Wang, M. et al. Bidirectional cross-kingdom RNAi and fungal uptake of external RNAs confer plant protection. Nature Plants 2 (2016). Artn 16151 10.1038/Nplants.2016.151

28 Weiberg, A. et al. Fungal Small RNAs Suppress Plant Immunity by Hijacking Host RNA Interference Pathways. Science 342, 118–123 (2013). 10.1126/science.1239705

29 Hudzik, C., Maguire, S., Guan, S., Held, J. & Axtell, M. J. Trans-species microRNA loci in the parasitic plant Cuscuta campestris have a U6-like snRNA promoter. The Plant Cell 35, 1834–1847 (2023). 10.1093/plcell/koad076

30 Johnson, N. R., dePamphilis, C. W. & Axtell, M. J. Compensatory sequence variation between trans-species small RNAs and their target sites. eLife 8, e49750 (2019). 10.7554/eLife.49750

31 Shahid, S. et al. MicroRNAs from the parasitic plant Cuscuta campestris target host messenger RNAs. Nature 553, 82–85 (2018). 10.1038/nature25027

32 Zhang, Z. L. et al. Cross- kingdom RNA interference mediated by insect salivary microRNAs may suppress plant immunity. P Natl Acad Sci USA 121 (2024). ARTN e2318783121 10.1073/pnas.2318783121

33 Guo, H. Y. et al. Plant-Generated Artificial Small RNAs Mediated Aphid Resistance. Plos One 9 (2014). ARTN e97410 10.1371/journal.pone.0097410

34 Prado, E. & Tjallingii, W. F. Effects of previous plant infestation on sieve element acceptance by two aphids. Entomologia Experimentalis Et Applicata 82, 189–200 (1997). 10.1046/j.1570-7458.1997.00130.x

35 Langmead, B., Trapnell, C., Pop, M. & Salzberg, S. L. Ultrafast and memory-efficient alignment of short DNA sequences to the human genome. Genome Biol 10, R25 (2009). 10.1186/gb-2009-10-3-r25

36 Dobin, A. et al. STAR: ultrafast universal RNA-seq aligner. Bioinformatics 29, 15–21 (2013). 10.1093/bioinformatics/bts635

37 Love, M. I., Huber, W. & Anders, S. Moderated estimation of fold change and dispersion for RNA-seq data with DESeq2. Genome Biol 15, 550 (2014). 10.1186/s13059-014-0550-8

38 Annacondia, M. L. et al. Aphid feeding induces the relaxation of epigenetic control and the associated regulation of the defense response in. New Phytologist 230, 1185–1200 (2021). 10.1111/nph.17226

39 Cedden, D. & Güney, G. Small RNAs in insects: emerging classes and functions. Curr Opin Insect Sci 73 (2026). ARTN 101458 10.1016/j.cois.2025.101458

40 Baumberger, N. & Baulcombe, D. C. Arabidopsis ARGONAUTE1 is an RNA Slicer that selectively recruits microRNAs and short interfering RNAs. Proc Natl Acad Sci U S A 102, 11928–11933 (2005). 10.1073/pnas.0505461102

41 Rajeswaran, R. et al. Sequencing of RDR6-dependent double-stranded RNAs reveals novel features of plant siRNA biogenesis. Nucleic Acids Research 40, 6241–6254 (2012). 10.1093/nar/gks242

42 Li, Z. C. et al. Origin, evolution and diversification of plant ARGONAUTE proteins. Plant Journal 109, 1086–1097 (2022). 10.1111/tpj.15615

43 Carbonell, A. Plant ARGONAUTEs: Features, Functions, and Unknowns. Plant Argonaute Proteins 1640, 1–21 (2017). 10.1007/978-1-4939-7165-7_1

44 Annacondia, M. L. & Martinez, G. Reprogramming of RNA silencing triggered by infection in Arabidopsis. Genome Biology 22 (2021). ARTN 340 10.1186/s13059-021-02564-z

45 Harvey, J. J. et al. An antiviral defense role of AGO2 in plants. PLoS One 6, e14639 (2011). 10.1371/journal.pone.0014639

46 Thody, J. et al. PAREsnip2: a tool for high-throughput prediction of small RNA targets from degradome sequencing data using configurable targeting rules. Nucleic Acids Res 46, 8730–8739 (2018). 10.1093/nar/gky609

47 He, S. et al. The LSD1-Interacting Protein GILP Is a LITAF Domain Protein That Negatively Regulates Hypersensitive Cell Death in Arabidopsis. PLOS ONE 6, e18750 (2011). 10.1371/journal.pone.0018750

48 Jing, Y., Guo, Q. & Lin, R. The SNL-HDA19 histone deacetylase complex antagonizes HY5 activity to repress photomorphogenesis in Arabidopsis. New Phytologist 229, 3221–3236 (2021). 10.1111/nph.17114

49 Song, C. P. et al. Role of an Arabidopsis AP2/EREBP-type transcriptional repressor in abscisic acid and drought stress responses. Plant Cell 17, 2384–2396 (2005). 10.1105/tpc.105.033043

50 Cabreira-Cagliari, C. et al. GILP family: a stress-responsive group of plant proteins containing a LITAF motif. Funct Integr Genomics 18, 55–66 (2018). 10.1007/s10142-017-0574-8

51 Szabó, A. K. et al. Local Aphid Species Infestation on Invasive Weeds Affects Virus Infection of Nearest Crops Under Different Management Systems - A Preliminary Study. Front Plant Sci 11, 684 (2020). 10.3389/fpls.2020.00684

52 Bown, D. P., Wilkinson, H. S. & Gatehouse, J. A. Differentially regulated inhibitor-sensitive and insensitive protease genes from the phytophagous insect pest, Helicoverpa armigera, are members of complex multigene families. Insect Biochem Mol Biol 27, 625–638 (1997). 10.1016/s0965-1748(97)00043-x

53 Guo, Y., Ji, N., Bai, L., Ma, J. & Li, Z. Aphid Viruses: A Brief View of a Long History. Front Insect Sci 2, 846716 (2022). 10.3389/finsc.2022.846716

54 Ng, J. C. & Perry, K. L. Transmission of plant viruses by aphid vectors. Mol Plant Pathol 5, 505–511 (2004). 10.1111/j.1364-3703.2004.00240.x

55 van Munster, M. et al. Characterization of a new densovirus infecting the green peach aphid Myzus persicae. J Invertebr Pathol 84, 6–14 (2003). 10.1016/s0022-2011(03)00013-2

56 Ravet, A. et al. Vesicular and non-vesicular extracellular small RNAs direct gene silencing in a plant-interacting bacterium. Nature communications 16, 3533 (2025). 10.1038/s41467-025-57908-1

57 Koch, B. L. et al. Arabidopsis Produces Distinct Subpopulations of Extracellular Vesicles That Respond Differentially to Biotic Stress, Altering Growth and Infectivity of a Fungal Pathogen. J Extracell Vesicles 14, e70090 (2025). 10.1002/jev2.70090

58 Koch, B. L., Singla-Rastogi, M. & Innes, R. W. Extracellular Vesicles and Extracellular RNAs in Plant-Microbe Interactions. Annu Rev Plant Biol 77, 225–255 (2026). 10.1146/annurev-arplant-063025-110704

59 Yu, X., Willmann, M. R., Anderson, S. J. & Gregory, B. D. Genome-Wide Mapping of Uncapped and Cleaved Transcripts Reveals a Role for the Nuclear mRNA Cap-Binding Complex in Cotranslational RNA Decay in Arabidopsis. Plant Cell 28, 2385–2397 (2016). 10.1105/tpc.16.00456

60 Menuet, K. et al. Aphid Salivary MIF Modulates Plant Programmed Cell Death and DNA Damage Response and Interacts with SOG1. bioRxiv, 2026.2004.2001.715815 (2026). 10.64898/2026.04.01.715815

61 Gravino, M. et al. Aphid effector Mp10 balances immune suppression and defence activation through EDS1-dependent modulation of plant DAMP responses. New Phytologist 248, 913–935 (2025). 10.1111/nph.70419

