## Supplementary figures and images for "Trans-kingdom delivery of aphid-derived small RNAs into *Arabidopsis thaliana* modulates plant immunity"

### Supplementary Figure 1

#### Supplementary Figure 1.

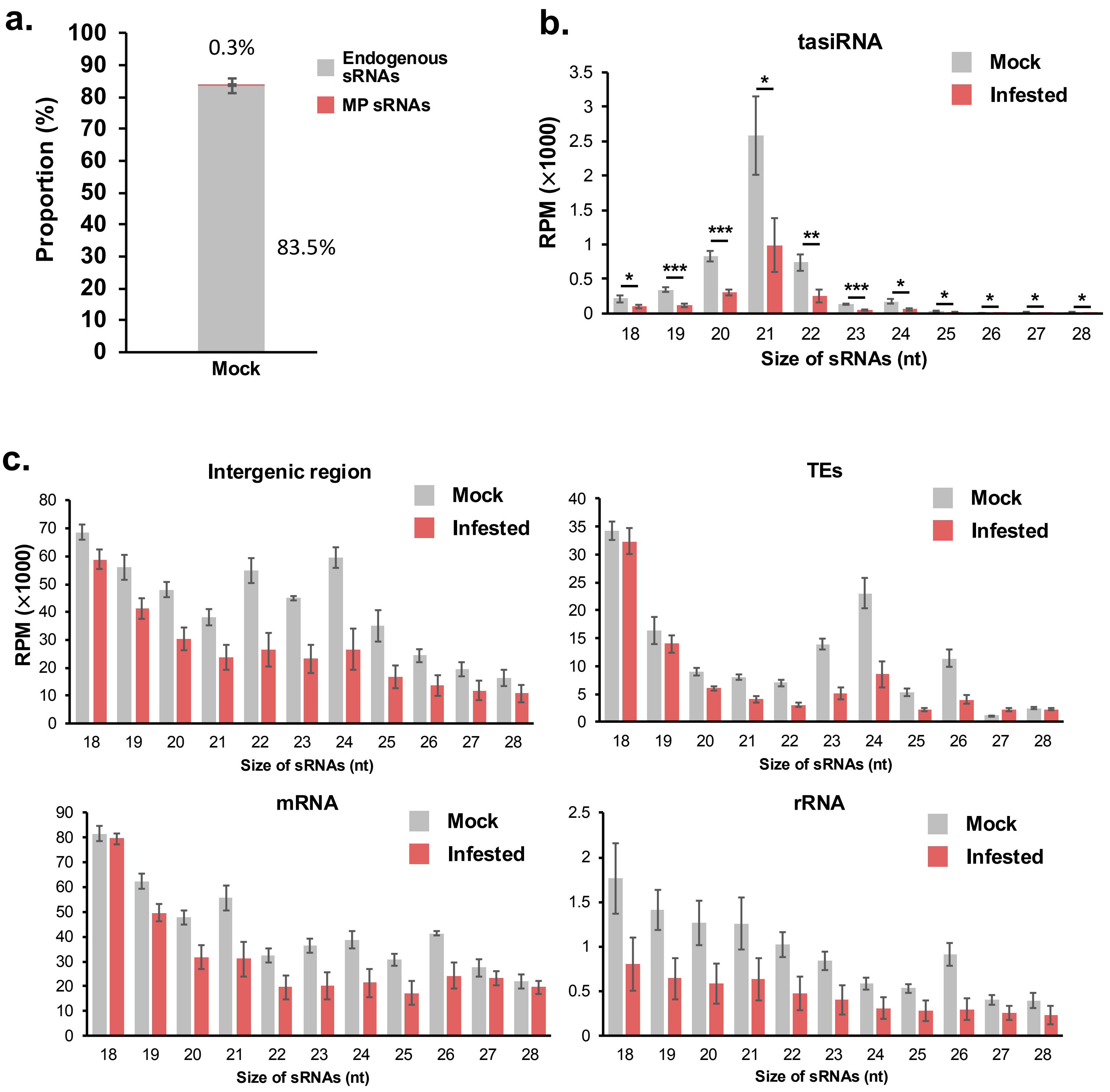

### Supplementary Figure 2

#### Supplementary Figure 2.

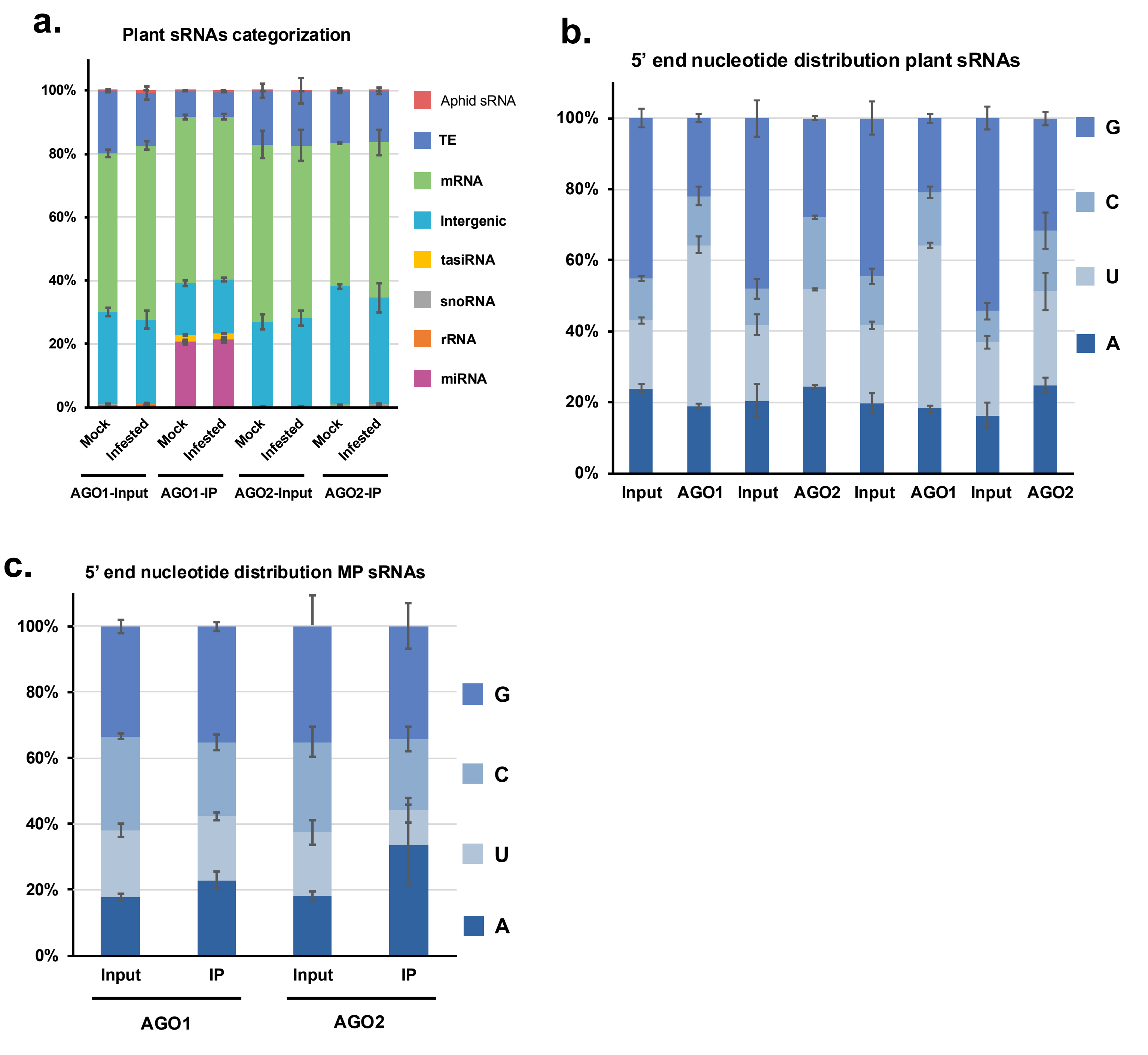

### Supplementary Figure 3

#### Supplementary Figure 3.

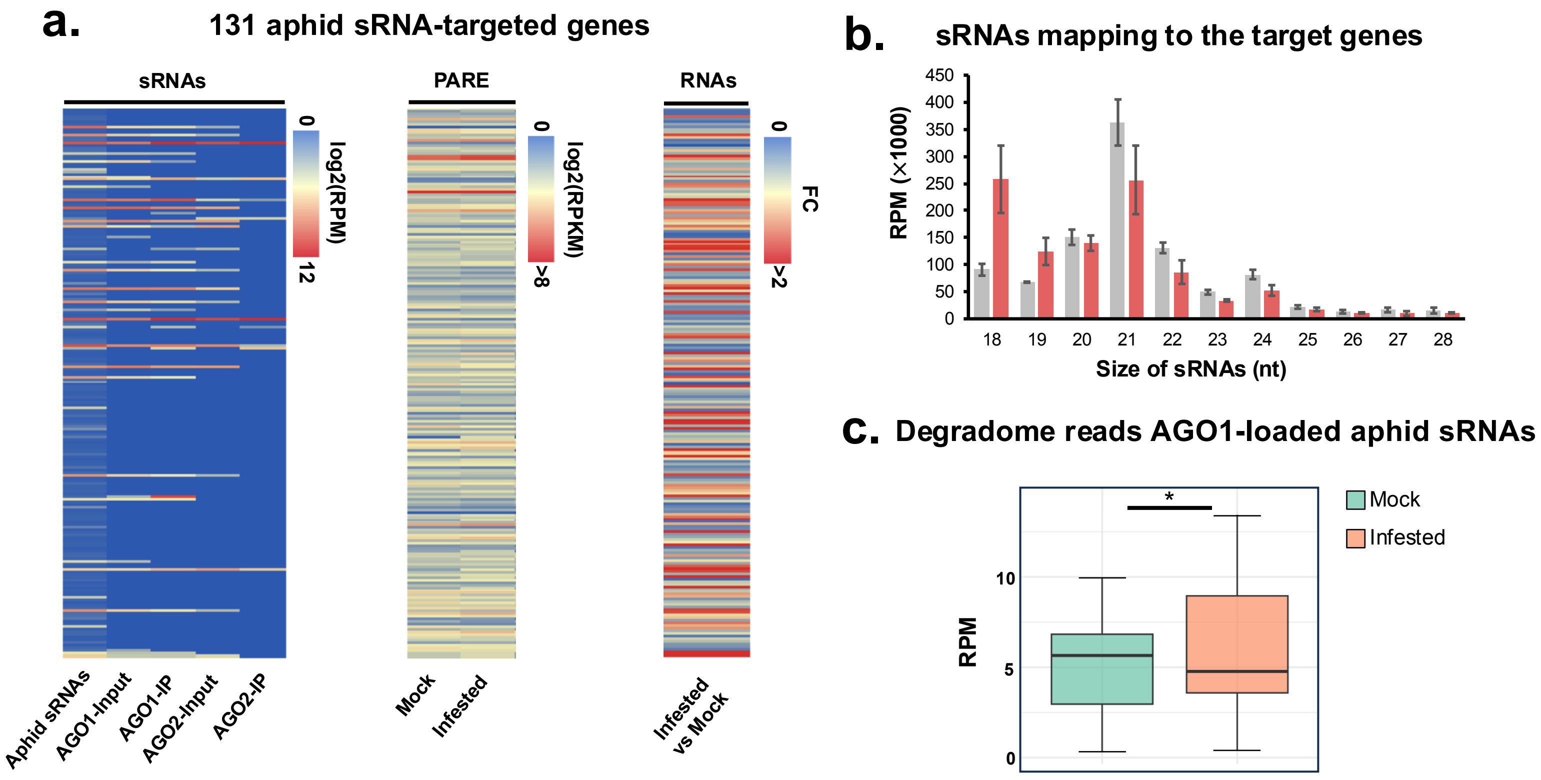

### Supplementary Figure 4

#### Supplementary Figure 4.

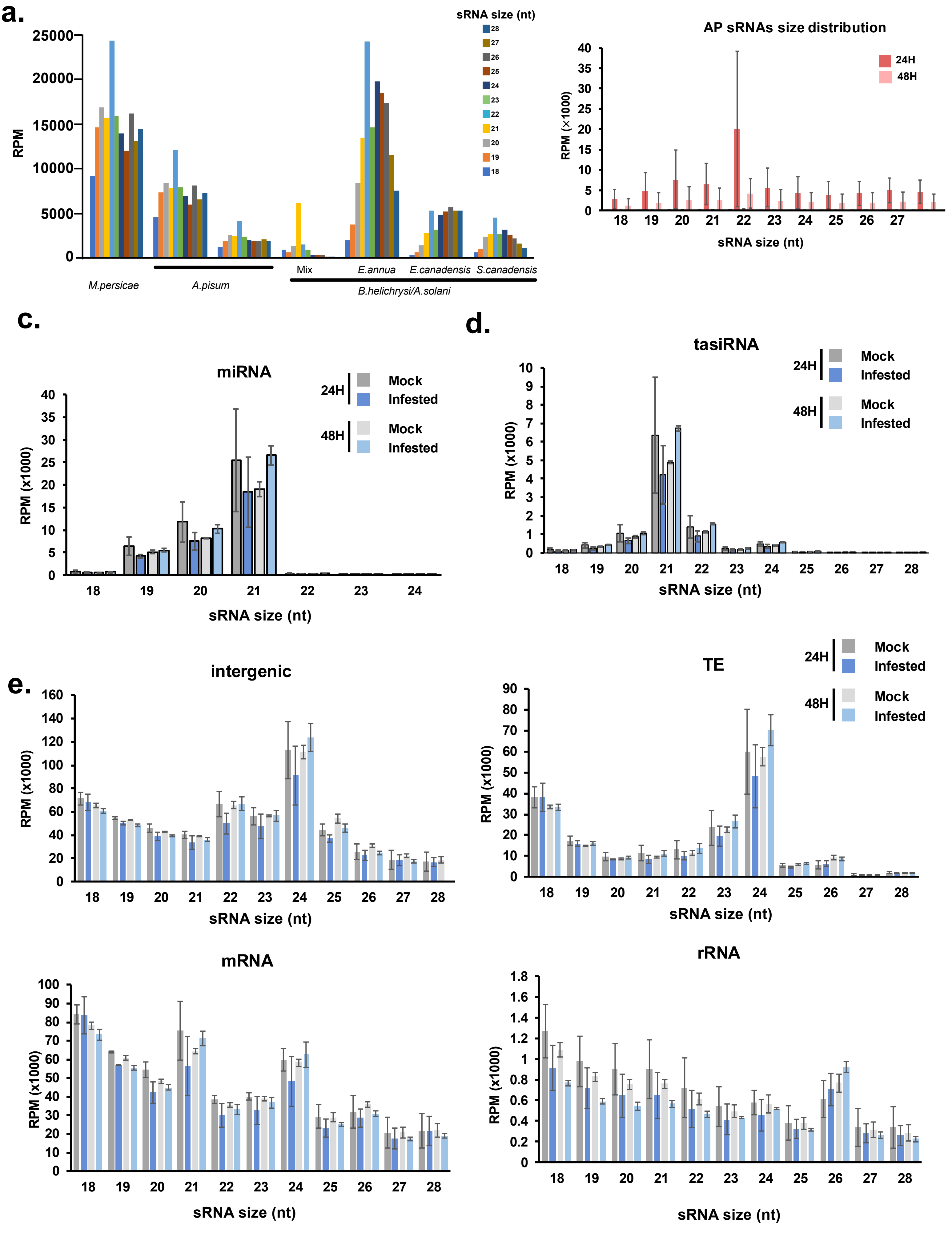

### Supplementary Figure 5

#### Supplementary Figure 5.

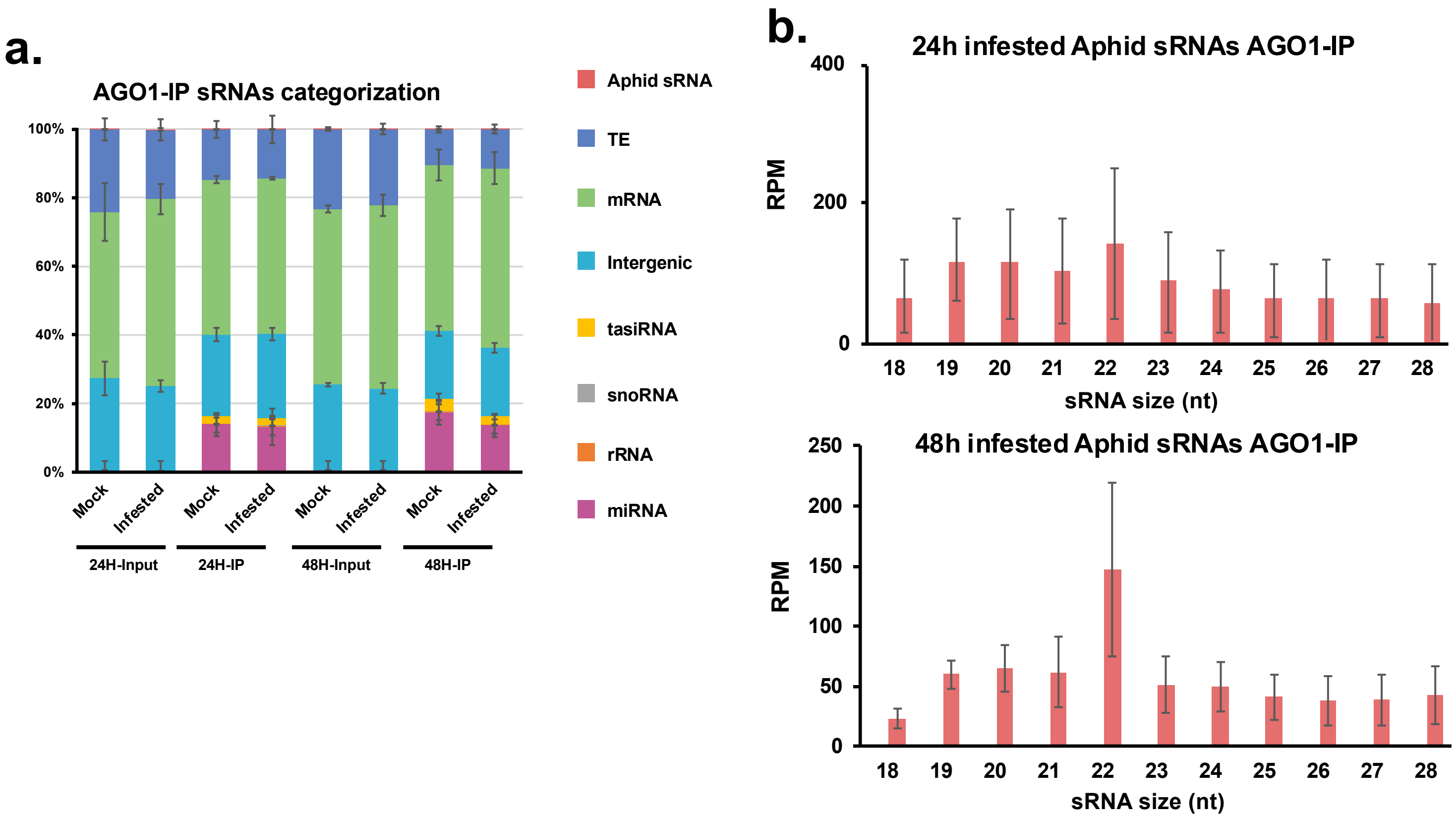
